# Finimizers: Variable-length bounded-frequency minimizers for *k*-mer sets

**DOI:** 10.1101/2024.02.19.580943

**Authors:** Jarno N. Alanko, Elena Biagi, Simon J. Puglisi

**Affiliations:** Helsinki Institute for Information Technology (HIIT) Department of Computer Science, University of Helsinki, Finland

**Keywords:** Minimizer, Finimizer, Spectral Burrows-Wheeler transform, pseudoalignment, *k*-mer, de Bruijn graph

## Abstract

The minimizer of a *k*-mer is the smallest *m*-mer inside the *k*-mer according to some order relation *<* of the *m*-mers. Minimizers are often used as keys in hash tables in indexing tasks in metagenomics and pangenomics. The main weakness of minimizer-based indexing is the possibility of very frequently occurring minimzers, which can slow query times down significantly. Popular minimizer alignment tools employ various and often wild heuristics as workarounds, typically by ignoring frequent minimizers or blacklisting commonly occurring patterns, to the detriment of other metrics (e.g., alignment recall, space usage, or code complexity).

In this paper, we introduce *frequency-bounded minimizers*, which we call *finimizers*, for indexing sets of *k*-mers. The idea is to use an order relation *<* for minimizer comparison that depends on the frequency of the minimizers within the indexed *k*-mers. With finimizers, the length *m* of the *m*-mers is not fixed, but is allowed to vary depending on the context, so that the length can increase to bring the frequency down below a user-specified threshold *t*. Setting a maximum frequency solves the issue of very frequent minimizers and gives us a worst-case guarantee for the query time. We show how to implement a particular finimizer scheme efficiently using the Spectral Burrows-Wheeler Transform (SBWT) (Alanko et al., Proc. SIAM ACDA, 2023) augmented with longest common suffix information. In experiments, we explore in detail the special case in which we set *t* = 1. This choice simplifies the index structure and makes the scheme completely parameter-free apart from the choice of *k*. A prototype implementation of this scheme exhibits *k*-mer localization times close to, and often faster than, stateof-the-art minimizer-based schemes. The code is available at https://github.com/ElenaBiagi/Finito.

## 1. Introduction

IN recent years, use of *minimizers* [9], [35], [36], [40] has become widespread in bioinformatics, particularly as an algorithmic tool to help manage large datasets for genome assembly [14] and for indexing tasks in pangenomics and metagenomics [19], [25]–[27], [33]. The minimizer of a *k*-mer is the smallest *m*-mer inside the *k*-mer. The scheme is parameterized by the integers *m* ≤ *k* and an order relation *<* of the minimizers. Typically, *k* ≈ 30, *m* ≈18 and *<* is either the lexicographic order, or an order determined by the hash values of the *k*-mers under some hash function.

In the context of indexing, a typical approach is to store location information for each minimizer of the data – for example, the coordinates of their occurrences in the sequences being indexed, which may be, e.g., reads, assembled genomes, or unitigs^1^ derived from such sequences. The index then consists of a mapping of each minimizer to a list of their occurrences, along with the indexed sequences themselves. With this information, it is possible to align query sequences of any length against the indexed data by computing the minimizers of the query sequence, locating them using the index, and then checking whether the sequence around the minimizers matches (or approximately matches) the query.

The main weakness of this indexing strategy is that the query time is proportional to the number of occurrences of the query’s minimizers in the indexed data, which can be very large. For example, in the unitigs of the human genome, minimizers can occur up to 3.6 × 10^4^ times for *k* = 31 and *m* = 20 [33]. Different tools attempt to get around this problem in different ways. One possible method is to use maximal exact matches as alignment seeds instead [24], but this comes with its own drawbacks regarding performance and sensitivity. The most popular minimizer-based alignment tool, MINIMAP2 [25], sidesteps the issue of frequent minimizers by ignoring minimizers that occur more often than a user-defined threshold. The SSHASH compact index for *k*-mer lookup [33] has a fallback for frequent minimizers, which can add a significant amount of space to the basic index. Nyström-Persson et al. [30] observe that handling of minimizers in *k*-mer counting software can be particularly involved:

“The *k*-mer counter KMC2 [12] introduced minimizer signatures, which order *m*-mers lexicographically, except that in order to reduce data skew, *m*-mers starting with AAA or ACA are given lower priority, and *m*-mers containing AA anywhere are also avoided, except for AA at the start … [t]his ordering is also used by Gerbil [15] and by FastKmer [17] in a modified form *with some additional rules*^2^”.

All these workarounds can have a detrimental effect on other important metrics, such as alignment recall, index size, and code complexity. *Another general difficulty with the use of minimizers is that the minimizer length m needs to be fixed before indexing*, even though it may be hard for the user to know what an efficacious value of *m* would be in advance.

In this paper, we introduce *frequency-bounded minimizers*, which we call *finimizers*. The idea is to use an order relation *<* for minimizer comparison that depends on the frequency of the minimizers themselves. With finimizers, the length *m* of the *m*-mers is not fixed, but is allowed to vary, so that the length can increase to bring the frequency down below a userdefined maximum threshold *t*. Setting a maximum frequency this way solves the issue of very frequent minimizers and gives us a worst-case guarantee for the query time.

Our specific context throughout is the indexing of unitigs (or, more generally, any disjoint spectrum preserving string set, *DSPSS*). In this context, a set of finimizers is guaranteed to exist: when the input is a *DSPSS*, all *k*-mers occur only once, so there are always low-frequency *m*-mers.

The principal advantages of finimizers for indexing are their bounded worst-case localization time and that they elide the need for the user to fix the minimizer length *m*. While it may appear we are exchanging one parameter for another (i.e., minimizer length for finimizer frequency), we show that it is possible to efficiently find sets of finimizers with frequency *t* = 1, making some specific manifestations of finimizers essentially parameter free.

Indeed, the main contribution of this article is to define and explore such a finimizer scheme—*Shortest Unique Finimizers*—that is fast to compute, compact to store, and can outperform state-of-the-art minimizer-based schemes when applied to *k*-mer localization. In practice, our prototype implementation of shortest-unique finimizers exhibits a *k*-mer localization index that takes 25-30 bits per *k*-mer on a wide range of datasets, making it only marginally larger than SSHASH. Simultaneously, it allows localization queries at comparable speeds with *asymptotic guarantees on performance* and *without the need to set parameters at construction time*.

At face value, finimizers may seem to come with their own set of drawbacks. Firstly, unlike with minimizers, computation is no longer local, as an efficient means of determining the frequency of a (potential) finimizer in the whole dataset is required. Moreover, the fixed-length nature of minimizers means they are particularly amenable to localization via slidingwindow hashing schemes: the potential minimizer at a given position is always the substring of length *m* starting there. With their variable length, finimizers afford no such luxury. However, we show how both these problems can be elegantly and scalably solved via the use of the Spectral BurrowsWheeler transform (SBWT) [4] in lieu of hashing. As a bonus, we also describe a fast heuristic for *unitig orientation* that can significantly reduce the size of SBWT-based indexes — indeed, by 25% or more in our experiments.

Apart from efficient *k*-mer localization queries, we envisage a vast array of possible future applications for finimizers. We list some of these ideas at the end of the paper.

### Related Work

Finimizers are tangentially related to, though decided different from, frequency-aware minimizer schemes [9], [20]. Chikhi et al. [9] suggest ordering minimizer candidates by frequency. In the robust winnowing scheme used by winnowmap [20] the set of minimizers occurring more than 1024 times is stored and those *m*-mers are assigned a lower weight during minimizer selection. Another superficially related technique is universal hitting sets, where for integers *k* and *L > k*, a set of *k*-mers is a universal hitting set if every possible *L*-length sequence must contain a *k*-mer from the set [32]. We emphasise that, unlike finimizers, these approaches still use fixed-length *m*-mers and provide no guarantees on minimizer frequencies.

In the context of *k*-mer localization queries, the state-of-theart in minimizer-based schemes is the recent and fast SSHASH of Pibiri [33]. To give SSHASH its due, it does offer asymptotic guarantees on localization time; however it does so by making a special case of frequent minimizers, an approach that does not always pan out well. For example, on a set of Nanopore E.coli reads^3^, setting minimizer length *m* = 14 increases relative index size by almost 30% (from 12.7 to 16.2 bits/kmer) even though *k decreases* from 31 to 21.

### Roadmap

The next section lays down notation and basic concepts used throughout. Section III formally defines finimizers, including the shortest unique finimizer scheme. Section IV then describes how to compute and query shortest unique finimizers as well as other details of our *k*-mer localization index. In Section V, we describe a heuristic for orienting unitigs that can significantly reduce SBWT-index size. In Section VI, we report statistics on finimizer length and density, and the results of a localization query benchmark. Conclusions, reflections and potential future directions are then offered in Section VII.

## II. Preliminaries

Throughout this paper, a *string X*[1..*n*] is a sequence of |*X*| symbols over the DNA alphabet Σ = { A, C, G, T}. The empty string is denoted *ϵ* and |*ϵ*| = 0. The *substring* of *X* starting at symbol *i* and ending at symbol *j* is denoted *X*[*i*..*j*]. We also use the half-open interval notations *X*(*i*..*j*] = *X*[*i* + 1..*j*] and *X*[*i*..*j*) = *X*[*i*..*j* −1]. A *prefix* is a substring with starting position 1 and a *suffix* is a substring with ending position *n*. A *k-mer* refers to a (sub)string of length *k*. A minimizer is the smallest *m*-mer of *k*-mer:

### Definition 1

*(Minimizer) Let Y be a k-mer, and m be an integer such that m* ≤ *k. The minimizer of Y is the smallest m-mer of Y according to a given order relation of minimizers*.

This order relation could be the simple lexicographic order or something more complex. The set of *k*-mers of a string *X* is called the *k-spectrum* of *X*.

### Definition 2

*(k-Spectrum) The k-spectrum S*_*k*_(*X*) *of string a X is the set of distinct k-mers* {*X*[*i*..*i* + *k* − 1] | *i* = 1, …, |*X*| − *k* + 1} *The k-spectrum S*_*k*_(*X*_1_, …, *X*_*m*_) *of a set of strings X*_1_, …, *X*_*m*_ *is the union* 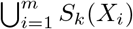

A *padded k-spectrum* adds a minimal set of $-padded *dummy k*-mers to ensure that in the de Bruijn graph (to be defined shortly), every non-dummy *k*-mer has an incoming path of length at least *k*:

### Definition 3

*(Padded k-Spectrum) Let R* = *S*_*k*_(*X*_1_, …, *X*_*m*_) *be a k-spectrum with alphabet* Σ, *and let R*^*′*^ ⊆ *S be the set of k-mers Y such that Y* [1..*k* − 1] *is not a suffix of any k-mer in R. The padded k-spectrum is the set* 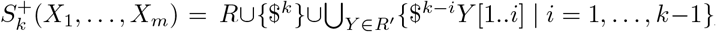, *with special character* $ ∉ Σ *and* $ *< c for all c* ∈ Σ.

For example, if *X*_1_ = ACGT, *X*_2_ = GACG and *k* = 3, then *S*_3_(*X*_1_, *X*_2_) = {ACG, CGT, GAC} and 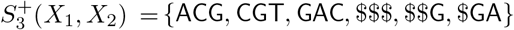. See Figure 3 for a more elaborate example.

The *de Bruijn graph* is an important concept in *k*-mer indexing. In this paper, we use a padded node-centric definition, where the de Bruijn graph of a set of strings *X*_1_, …, *X*_*m*_ is an edge-labeled graph *G* = (*V, E*), where 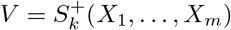 and there is an edge from *k*-mer *u* to *υ* iff *u*(1..*k*] = *υ* [1..*k*). See Figure 1 for an example. The label of an edge *e* = (*u, υ*), denoted with *𝓁*(*e*) or *𝓁*(*u, υ*), is the character *υ* [*k*] (with this, $ never appears as an edge label). The label of a path of edges is the concatenation of the edge labels in the order of the path, and the label of a path of nodes *υ*_1_, …, *υ*_*m*_ is *υ*_1_ ·*𝓁*(*υ*_1_, *υ*_2_) … *𝓁*(*υ*_*m*−1_, *υ*_*m*_), with possible dollar symbols in *υ*_1_ removed.

**Fig. 1.**
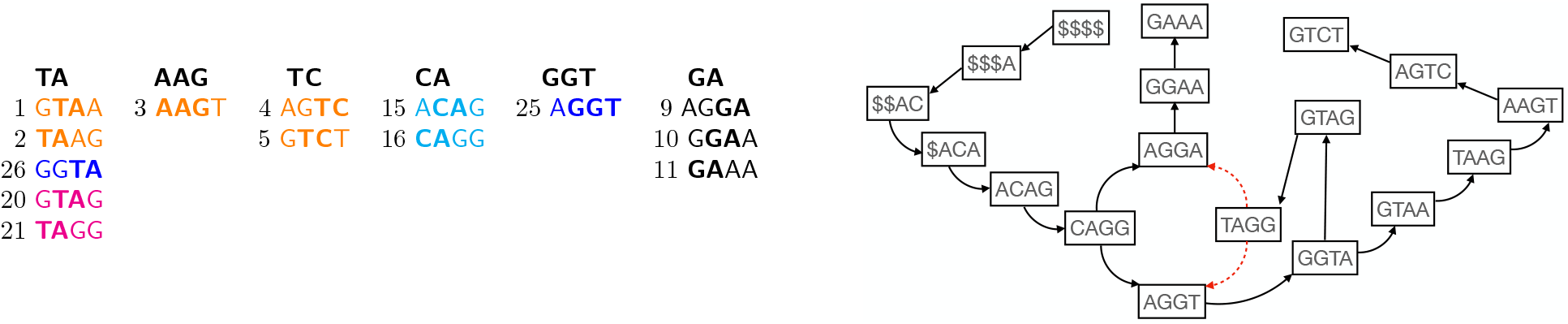
Left: A set of shortest unique finimizers of the set of unitigs *U* = {GTAAGTCT, AGGAAA, ACAGG, GTAGG, AGGTA}, *k* = 4. From the *SFS*_*k*,1_ of each unitig, we can obtain the set of shortest unique finimizers (*SUF*s) for each *k*-mer. *SFS*_*k*,1_ = [⊥, ⊥, (2, 1, 8), (3, 1, 5), (3, 1, 11), (3, 1, 17), (2, 1, 10), (2, 1, 16)], [⊥, ⊥, ⊥, (2, 1, 7), (3, 1, 4), (3, 1, 3)], [⊥, (2, 1, 9), (2, 1, 6), (3, 1, 12), (4, 1, 14)], [⊥, ⊥, (2, 1, 8), (3, 1, 13), (4, 1, 15)], [⊥, ⊥, ⊥, (3, 1, 18), (2, 1, 8)]. These five strings, composed of 14 *k*-mers, can be represented with six finimizers. For the *k*-mer ACAG we choose CA over AC as CA*<*_*colex*_AC. The *SUF* for the *k*-mer GGTA is TA, since it is shorter than GGT. Right: The de Bruijn graph of *U*. Red dashed edges are pruned from the graph because nodes they point to can be reached from other (black) edges.

A *unitig* is a maximal simple node path *υ*_1_, …, *υ*_*m*_ in a de Bruijn graph such that outdegree(*υ*_*i*_) = 1 for *i >* 1 and indegree(*υ*_*i*_) = 1 if *i < m*. Maximality in this context means that the path can not be extended in either direction without repeating nodes or breaking indegree or outdegree constraints. The set of all unitigs covers all nodes of the graph and is uniquely defined if a canonical choice of starting point is made for disjoint cycles (for example, the lexicographic minimum).

### Definition 4

*(Disjoint Spectrum-Preserving String Set (DSPSS)) A set of strings X*_1_, …, *X*_*m*_ *is a disjoint spectrumpreserving string set of order k of a set of strings Y*_1_, … *Y*_*n*_ *iff S*_*k*_(*X*_1_, …, *X*_*m*_) = *S*_*k*_(*Y*_1_, …, *Y*_*n*_) *and all k-mers in X*_1_, …, *X*_*m*_ *are distinct*.

The set of node path labels of all unitigs in the de Bruijn graph of a set of strings in an example of a *DSPSS* for that set of strings [34], and the smallest *DSPSS* can be computed in linear time using the Eulertigs algorithm [38].

The main operation we are interested in supporting on a *k*-spectrum is the *localization* query:

### Definition 5

*(k-mer localization) Given a disjoint spectrumpreserving string set 𝒟 and k-mer Y, localize(Y) returns either* ⊥*if the k-mer does not occur in any X* ∈ 𝒟, *or otherwise a pair* (*i, j*), *where i is the index of the string X*_*i*_ ∈ 𝒟 *that contains Y, and j is the starting point of Y in X*_*i*_.

Given a DSPSS 𝒟= *X*_1_, …, *X*_*m*_, it will be important to make a distinction between DSPSS occurrences and DBG occurrences of substrings:

### Definition 6

*(DSPSS occurrence). Given a DSPSS* 𝒟= *X*_1_, …, *X*_*m*_, *a DSPSS occurrence of string Y is a pair* (*i, j*) *such that X*_*i*_[*j*..*j* + |*Y* | − 1] = *Y*.

### Definition 7

*(DSPSS frequency) Given a DSPSS* 𝒟 = *X*_1_, …, *X*_*m*_, *the DSPSS frequency of Y is the number of distinct DSPSS-occurrences of Y*.

### Definition 8

*(DBG occurrence) Given a DSPSS* 𝒟 = *X*_1_, …, *X*_*m*_, *a DBG occurrence of Y is a node of the de Bruijn graph of* 𝒟 *that has an incoming path with edges spelling Y*.

### Definition 9

*(DBG frequency) Given a DSPSS* 𝒟 = *X*_1_, …, *X*_*m*_, *the DBG frequency of Y is the number of DBG occurrences of Y*.

Note that the number of times a substring occurs in the de Bruijn graph is *not* necessarily equal to the number of occurrences in the *DSPSS*, since, for example, the labels of neighboring unitigs in the unitig DSPSS overlap by *k* − 1 characters. This is illustrated in Figure 2, where the DSPSS frequency of CA is 5, but the DBG frequency is only 1. The DBG frequency equals the number of *k*-mers in the padded *k*-spectrum that have *Y* as a suffix.

**Fig. 2.**
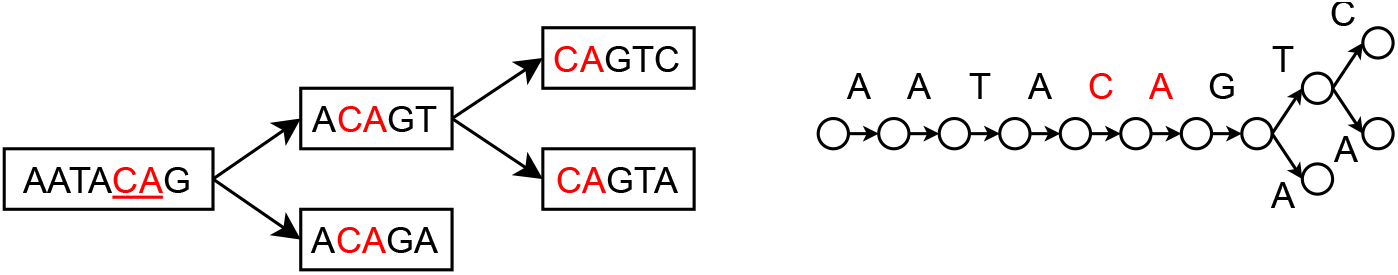
A finimizer that is unique in the de Bruijn graph (right, *k* = 5) may not be unique in the unitig DSPSS (left). In our index structure, the unitig occurrence that represents the de Bruijn graph occurrence is the one which is observed farthest from beginning of the unitig.

**Fig. 3.**
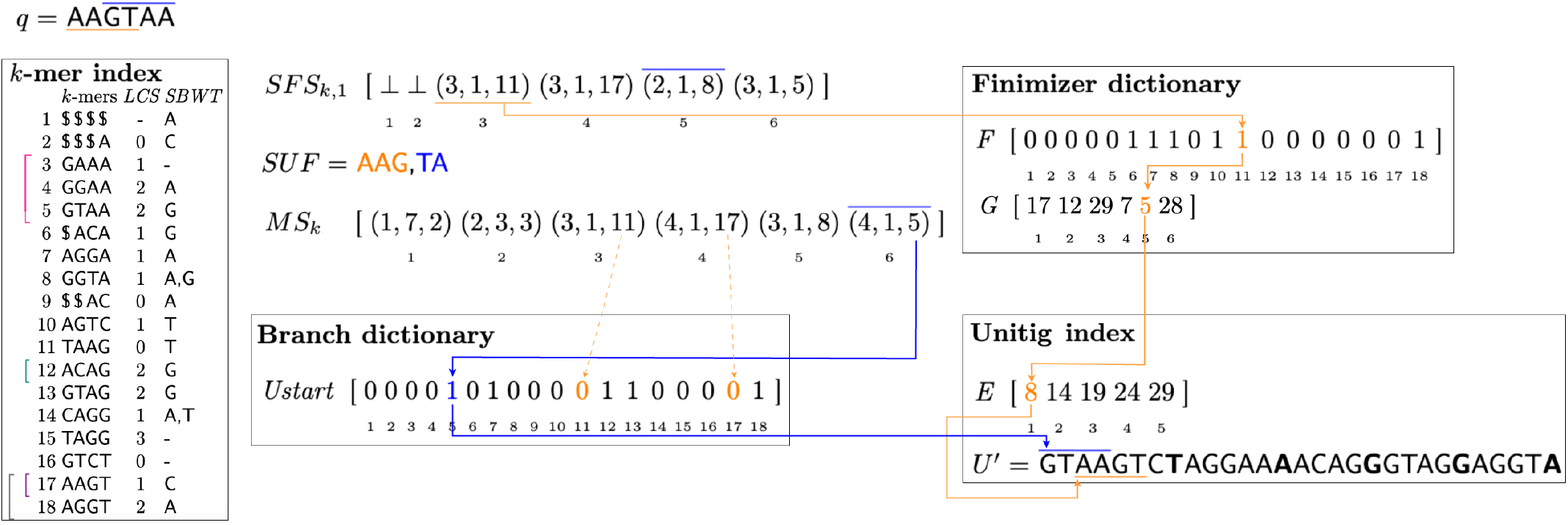
A schematic representation of the proposed data structure for localization queries on the running example. Given the *k-mer index* we compute for each position of the query, from left to right, the shortest unique suffixes array *SFS*_*k*,1_, which gives us the shortest unique finimizers *SUF*, and the *k*-bounded matching statistics *MS*_*k*_. To the left we illustrate the *ExtendRight* operation, with extending character G and (magenta) initial interval [3, 5]_AA_, which leads to the (teal) interval [11, 11]_AAG_, further extended to [17, 17]_AAGT_(violet). We can perform a single *ContractLeft* operation without modifying this interval. A further left contraction would lead to the (black) interval [17, 18]_GT_. The flow of the algorithm is represented with arrows. From *MS*_*k*_ we can identify the rightmost branch for each *k*-mer in the query, if it exists. The *k*-mer (orange, *i* = 4) AAGT is found in a non-branching path in the DBG, thus we have to look up the *finimizer dictionary*. The finimizer is AGG and its colexicographic interval in the *SBWT* is [11, 11]. The global offset of the finimizer is obtained with a rank query on *F, G*[*F*.*rank*(11)] = 5. We can get the unitig identifier with a predecessor query on *E* (*unitig index*), *E*.*pred*(5) = 1. The starting point of the *k*-mer is 3 = 5 − 0 + (4 − 3) − *k* + 1 Finally AAGT is localized in *U* ^*′*^ with result (1, 3). The localization query works differently for GTAA (blue, *i* = 6) as we can identify a branch, at index 6, thus we know that the *k*-mer appears at the beginning of a unitig, which is identified with a rank query on *Ustart, Ustart*.*rank*(5) = 1. The starting point of the *k*-mer is 1 = 6 − 6 + 1. This last *k*-mer of the query is then localized in *U* ^*′*^ with output (1, 1).

A path *υ*_1_, … *υ*_*n*_ in a graph is considered branching if there exists *i >* 1 such that the indegree of *υ*_*i*_ is at least 2, or there exists *i < n* such that the outdegree of *υ*_*i*_ is at least 2. A string *Y* is considered branching in a DBG if there exists a branching path labeled with *Y*.

A key tool in succinct data structures is support for *rank* and *select* queries on a bit string (or bit vector) *Z* of length *n*. For *i* ≤ *n* and *z* ∈ {0, 1}, *rank*_*z*_(*i*) is number of *z*’s in the first *i* bits of *Z*, and *select*_*z*_(*i*) is the position of the *i*th *z* in *Z*. Classical techniques need *n* + *o*(*n*) bits to support each operation in *O*(1) time [29].

## III. Finimizer Functions

### Definition 10

*(Finimizer function) Let R be a set of k-mers and t be a positive integer. A function f* : Σ^*k*^ → Σ^*^ *is a t-finimizer function for R if for all X* ∈ Σ^*k*^, *f* (*X*) *is a substring of X, and f* (*X*) *has at most t occurrences in the de Bruijn graph of R. The set* { *f* (*X*) | *X* ∈ *R*} *is called a t-finimizer set for R*.

A *t*-finimizer function always exists because the identity function fulfills the requirements: the number of occurrences of every full *k*-mer is 1, so the frequency requirements are fulfilled. For the purposes of indexing, however, we desire a finimizer function such that the size of the image { *f* (*X*) |*X* ∈ *R*} is small, and such that adjacent *k*-mers in the de Bruijn graph likely have the same finimizer. Additionally, the function should allow a compact representation that also ensures fast evaluation.

There are potentially many finimizer functions with these properties. The one we focus on for the remainder of this paper is the function that picks the *shortest* substring with frequency at most *t*.

### Definition 11

*(Shortest t-Finimizers) Let G be the de Bruijn graph of a set of k-mers R, t* ≥ 1 *be an integer, X be a k-mer, and Y be a substring of X. We say Y is a* shortest *t*-finimizer *of X with respect to the k-mer set R if Y has at most t occurrences in G and there does not exist a shorter substring of X with at most t occurrences in G*.

If multiple shortest finimizer candidates exist, a consistent tiebreaker such as colexicographic comparison can be used to always pick the same finimizer.

When *t* = 1, we call the finimizers *Shortest Unique Finimizers (SUFs)*. As we shall see in Section IV, when implemented with care, a finimizer function for shortest unique finimizers can be constructed in time linear in the total length of the input *DSPSS*, and evaluated for every *k*-mer of a query sequence *Q* in time *O*(|*Q*|).

## IV. Computing and Querying Shortest Unique Finimizers

In this section we show how to index a set of unitigs for *k*-mer localization queries using Shortest Unique Finimizers (*SUF*s). The problem is related to the active line of research on shortest unique substring queries [1]. Here, we present a solution specialized to sets of *k*-mers by using the Spectral Burrows-Wheeler transform [4]. The methods generalize to any *DSPSS*, but here we focus on unitigs for concreteness. We first show how the shortest unique finimizer for each *k*-mer in the *DSPSS* can be identified in linear time in the total length of the *DSPSS* using the Spectral Burrows-Wheeler transform (*SBWT*) augmented with the longest-common-suffix (*LCS*) array, two data structures that we define below. See Figure 3 for an illustration of these structures.

### Definition 12

*(Spectral Burrows-Wheeler Transform (SBWT) [4]) Let R*^+^ *be a padded k-spectrum and let X*_1_, …, *X*_|*R*|_ *be the elements of R*^+^ *in colexicographic order. The SBWT is the sequence of sets A*_1_, …, *A*_|*R*|_ *with A*_*i*_ ⊆ Σ *such that A*_*i*_ = ∅ *if i >* 1 *and X*_*i*_[2..*k*] = *X*_*i*−1_[2..*k*], *otherwise A*_*i*_ = {*c* ∈ Σ | *X*_*i*_[2..*k*]*c* ∈ *R*^+^}.

It is helpful to think of the *SBWT* as encoding the set *R* of *k*-mers as a list arranged in colexicographical order. In the context of *k*-mer lookup, the *SBWT* allows navigation of that ordered list via the operation *ExtendRight*, defined as follows.

Let [*s, e*]_*α*_ denote the *colexicographic interval* of string *α*, where *s* and *e* are respectively the colexicographic ranks of the smallest and largest *k*-mer in the *SBWT* having substring *α* as a suffix. The *right extension* of the interval [*s, e*]_*α*_ with a character *c* ∈ Σ, denoted *ExtendRight*([*s, e*]_*α*_, *c*), is the interval [*s*^*′*^, *e*^*′*^]_*αc*_, or ⊥ if no such interval exists. *ExtendRight* can be answered in *O*(1) time using a data structure of *n* |Σ| +*o*(*n* |Σ|) bits [4].

It is easy to see that an *existential k*-mer lookup algorithm, which tells if a query *k*-mer *r* exists (or not) in *S*, comes from the repeated application of *ExtendRight* to the symbols of the query *k*-mer, starting with the interval [1, *n*]_*ϵ*_, where *ϵ* is the empty string. Indeed the algorithm visits all intervals of the colexicographically sorted *k*-mer spectrum that correspond to all prefixes of the query *k*-mer that are suffixes of *k*-mers in *S*. This also gives us the DBG frequency of every prefix of the query *k*-mer since the DBG frequency of a string is equal to the length of its colexicographic interval. To support efficient streaming queries, we pair the *SBWT* with the *LCS* array:

### Definition 13

*(Longest Common Suffix (LCS) Array [2]) Let R*^+^ *be a colexicographically ordered padded k-spectrum and let X*_*i*_ *be the colexicographically i-th k-mer of R*^+^. *The LCS array is an array of length* |*R*^+^| *such that LCS*[1] = 0, *and for i >* 1, *LCS*[*i*] *is the length of the longest common suffix of X*_*i*−1_ *and X*_*i*_.

In the above definition, we consider the empty string as a common suffix of any two *k*-mers, such that the longest common suffix is well-defined for any pair of *k*-mers.

The *LCS* array can be constructed from the *SBWT* in *O*(*n*) time [2]. It allows a different navigation operation—called

*ContractLeft*—on the colexicographically-ordered padded *k*-mer spectrum. In particular, operation *ContractLeft*([*s, e*]_*α*_) for |*α*| *>* 0 returns the interval [*s*^*′*^, *e*^*′*^]_*α*[2..|*α*|]_, A *ContractLeft* query can be answered by setting *s*^*′*^ to be the largest position smaller or equal to *s* in *LCS*, such that *LCS*[*s*^*′*^] *<* |*α*| − 1 (a *previous-smaller-value* query, PSV-query for short), or to 1 if no such position exists. Symmetrically, *e*^*′*^ can be found with a next-smaller-value (NSV) query respect to *e* (see Figure 3). There exist data structures that can answer both PSV and NSV queries in constant time with modest space overhead [7], [18]. In practice however, we implement *ContractLeft*([*s, e*]_*α*_) by simply scanning the *LCS* array left from *s* and right from *e*. This works well because in our algorithms, in practice, the vast majority of scans tend to be very short (at most 6 elements on average in our experiments). The scanning solution is also memory-local and is thus likely faster than more sophisticated data structures with worst-case guarantees.

The *ContractLeft* operation in combination with the *ExtendRight* operation allows us to compute in linear time the *k-bounded matching statistics*:

### Definition 14

*(k-bounded matching statistics) The k-bounded matching statistics of a query sequence Q*[1..*m*], *versus a padded k-spectrum R*^+^, *is the array MS*_*k*_[1..*m*] *such that MS*_*k*_[*i*] = (*d*_*i*_, *p*_*i*_, *𝓁*_*i*_), *where d*_*i*_ *is the largest non-negative integer such that Q*(*i* − *d*_*i*_..*i*] *is a suffix of at least one k-mer in R*^+^, *p*_*i*_ *is the DBG frequency of Q*(*i* − *d*_*i*_..*i*] *and 𝓁*_*i*_ *is the start of the colexicographic interval of Q*(*i* − *d*_*i*_..*i*].

We can compute *MS*_*k*_ efficiently by using a *k*-bounded variant of the BWT-based unbounded matching statistics algorithm of Ohlebusch et al. [31]. The pseudocode is given in Algorithm 1, and the proof of correctness is given in the Supplement. We will use the *MS*_*k*_ array during localization queries to find the colexicographic ranks of all *k*-mers of the query present in the index: all the *k*-mers ending at positions where the match length is equal to *k*.

Building on the idea of Algorithm 1, we can also compute the *shortest frequency-bounded suffixes*:

### Definition 15

*(Shortest frequency-bounded suffixes) The shortest suffix statistics of a query sequence Q*[1..*m*], *versus a padded k-spectrum R*^+^, *is the array SFS*_*k,t*_[1..*m*] *such that SFS*_*k,t*_[*i*] = (*d*_*i*_, *p*_*i*_, *𝓁*_*i*_), *where d*_*i*_ *is the smallest positive integer such that Q*(*i* − *d*_*i*_..*i*] *is a suffix of p*_*i*_ *(p*_*i*_ ≤ *t) k-mers in R*^+^, *and 𝓁*_*i*_ *is the start of the colexicographic interval of Q*(*i*−*d*_*i*_..*i*]. *If no such d*_*i*_ *exists, SFS*_*k,t*_[*i*] = ⊥.

The *SFS*_*k,t*_ array will be useful for identifying the shortest frequency-bounded substring of every *k*-mer in a query. Computation of the *SFS*_*k,t*_ array with unbounded *k* has been studied by Belazzougui and Cunial [5]. Their algorithm uses data structures on the suffix tree topology of the data to implement left contractions. We avoid most of the complications in their algorithm by replacing the suffix tree topology with the *LCS* array, which uses only ⌊log(*k* −1) ⌋ +1 bits per *k*-mer since the *LCS* values are in the range [0, *k* −1]. Algorithm 2 gives the pseudocode in terms of left contractions and right extensions. We give a sketch of the proof of correctness here. The full details can be found in the supplement.

### Algorithm 1

*k*-bounded matching statistics.

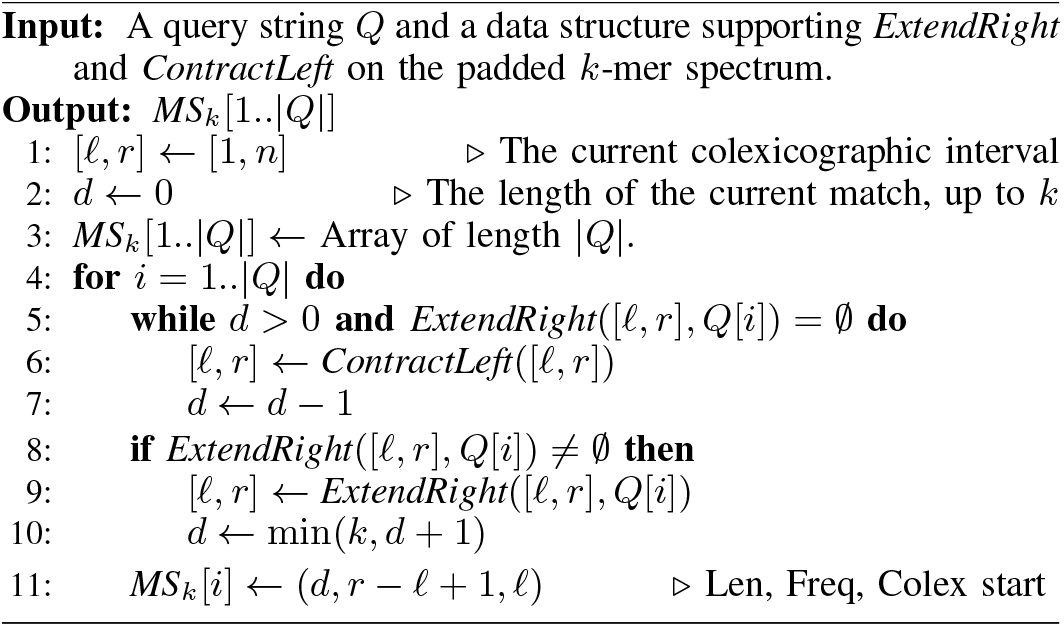

### Algorithm 2

Shortest frequency-bounded suffix statistics.

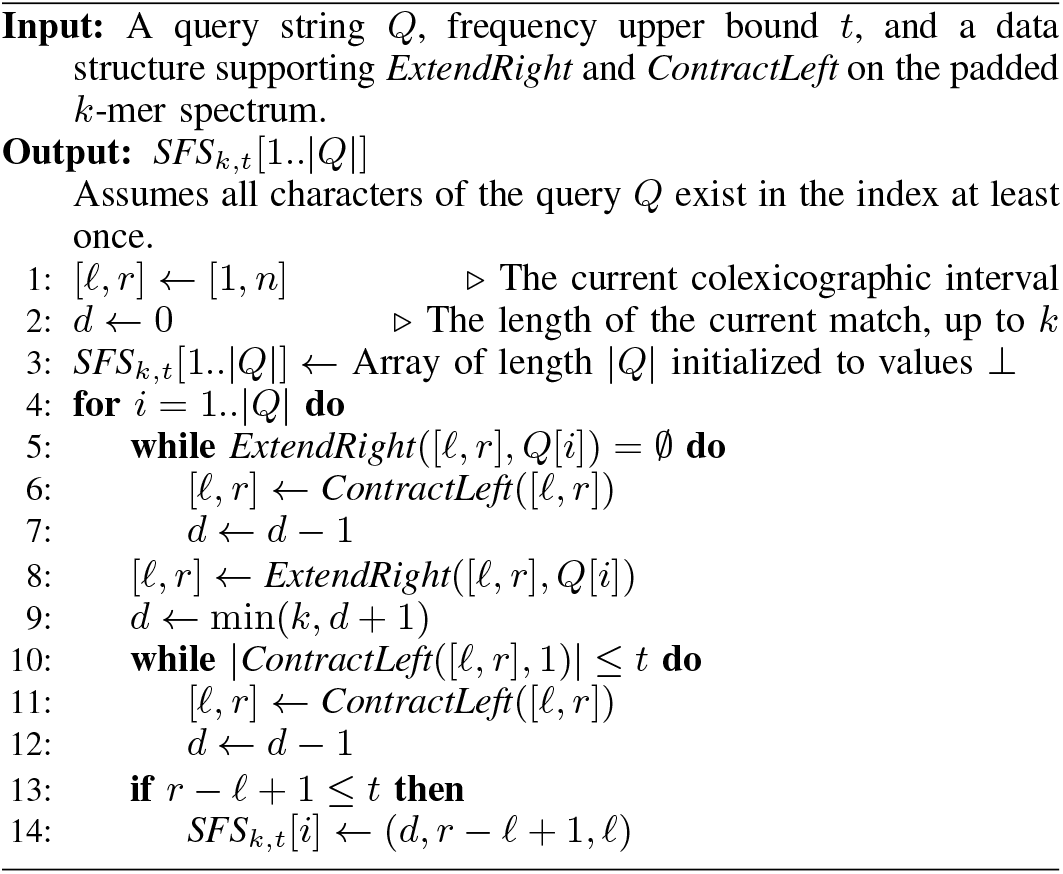

### Theorem 1

*Given constant-time ContractLeft and ExtendRight queries on padded k-spectrum R*^+^, *Algorithm 2 computes SFS*_*k,t*_ *for query Q in time O*(|*Q*|).

*Proof*. The algorithm maintains an invariant that at the end of iteration *i*, the interval [*𝓁, r*] is the interval of the shortest suffix of *Q*[1..*i*] occurring as a suffix of at most *t k*-mers, if it exists, or otherwise [*𝓁, r*] is the longest suffix of *Q*[1..*i*] occurring in the *k*-mers. Time is linear in |*Q*| since each query position is extended to and contracted from only once.

### A. Shortest Unique Finimizer Index Structure

Our index structure is composed of four main components. We describe these components in this subsection to give an overall view of our data structure. Subsequent sections show how they are constructed efficiently and finally how they are used for querying.

Let us denote the set of unitigs as *U* = {*u*_1_, *u*_2_, …, *u*_|*U*|_} sorted by the colexicographical order of their first *k*-mers. Let *n* denote the number of *k*-mers in the padded *k*-mer spectrum *R*^+^ of a set of *U*. It will be convenient for us to refer to *k*-mers in *R*^+^ by their colexicographical rank. Let *X*_*i*_ denote the *i*-th *k*-mer in *R*^+^ in colexicographic order. Our *SUF* index for *U* consists of the following components (see Figure 3 for an example):

1. The **k-mer index**. The *k*-mer index includes the *SBWT* and the *LCS* array, which are used to identify the shortest unique finimizers using *ExtendRight* and *ContractLeft* operations.

2. The **unitig index**. The purpose of this component is to provide random access to the unitig strings. It consists of the concatenation *U*^*′*^ = *u*_1_ ◦… ◦ *u*_*n*_, and the array *E* storing the endpoints of each unitig in *U*^*′*^ in ascending order, compressed with the Elias-Fano method (see [29]), with support for access and predecessor queries.

3. The **finimizer dictionary**. This component is used to map finimizers identified by their colexicographic ranks to their locations (“global offsets”) in *U*^*′*^. The finimizer dictionary stores a bit vector *F* [1..*n*] such that *F* [*i*] = 1 iff *X*_*i*_ has a *SUF* as a suffix. Note that a bit vector suffices for *F* precisely because the *SUF*s are unique w.r.t the de Bruijn graph. The bit vector *F* is processed for constant-time rank queries. The finimizer index also stores an array *G* such that *G*[*i*] is the ending position in *U*^*′*^ of an occurrence of the finimizer with colexicographic rank *i*. Note that even though finimizers are unique in the DBG, they may occur in multiple unitigs strings. To resolve this ambiguity, we always pick the uniquely defined unitig occurrence that occurs furthest from the start of a unitig. The correctness of the query algorithm will depend crucially on this choice.

4. The **branch dictionary**. The purpose of this component is to identify *k*-mers that are the first *k*-mers of their unitigs. To this end, we store a bit vector *UStart*[1..*n*] that marks the *k*-mers that are prefixes of unitigs. That is, *UStart*[*i*] = 1 iff the *X*_*i*_ is a prefix of some *u*_*j*_ ∈ *U*. The bit vector *UStart* is preprocessed for constant-time rank and select queries. This component will be important when localizing *k*-mers that cross branches in the DBG.

### B. Shortest Unique Finimizer Index Construction

In this section, we describe the construction of our index for shortest unique finimizers. We begin by sorting the unitigs by the colexicographic ranks of their first *k*-mers, and concatenating the sorted unitigs to construct *U*^*′*^ and *E*. Then, we construct the *SBWT* using the algorithm described in [4] and from there, build the *LCS* array using the techniques in [2]. The bit vector *Ustart* is easily constructed by searching the first *k*-mer of each unitig on the *SBWT* and marking the colexicographic positions of those.

Algorithm 3 gives pseudocode for constructing the finimizer dictionary, consisting of the global offsets array *G* and the finimizer colexicographic marks bit vector *F*. The algorithm processes the unitig strings one by one in the same order as they are concatenated in *U*^*′*^. For each unitig, we run Algorithm 2 to compute the shortest unique suffixes array, *SFS*_*k*,1_, (Def. 15) of the unitig. Then, we slide a window of length *k* over the *SFS*_*k*,1_ array while maintaining a monotonic queue data structure^4^ to be able to identify the shortest unique suffix inside the current *k*-mer window, with colexicographic tiebreaking. This process gives us the colexicographic rank and ending position of the finimizer of each *k*-mer of the unitig. We then mark these colexicographic ranks in the bit vector *F*, and store corresponding the global offsets. If a global offset is already stored for some colexicographic position, we overwrite it iff the finimizer ending position is further from the start of the unitig than the currently stored offset. The array *D* in the pseudocode keeps track of the distance of the stored global offset from the start of a unitig. For example, in Figure 2 the final global offset stored for CA would be the one that is contained in the leftmost unitig of the picture.

#### Algorithm 3

Shortest Unique Finimizers dictionary construction

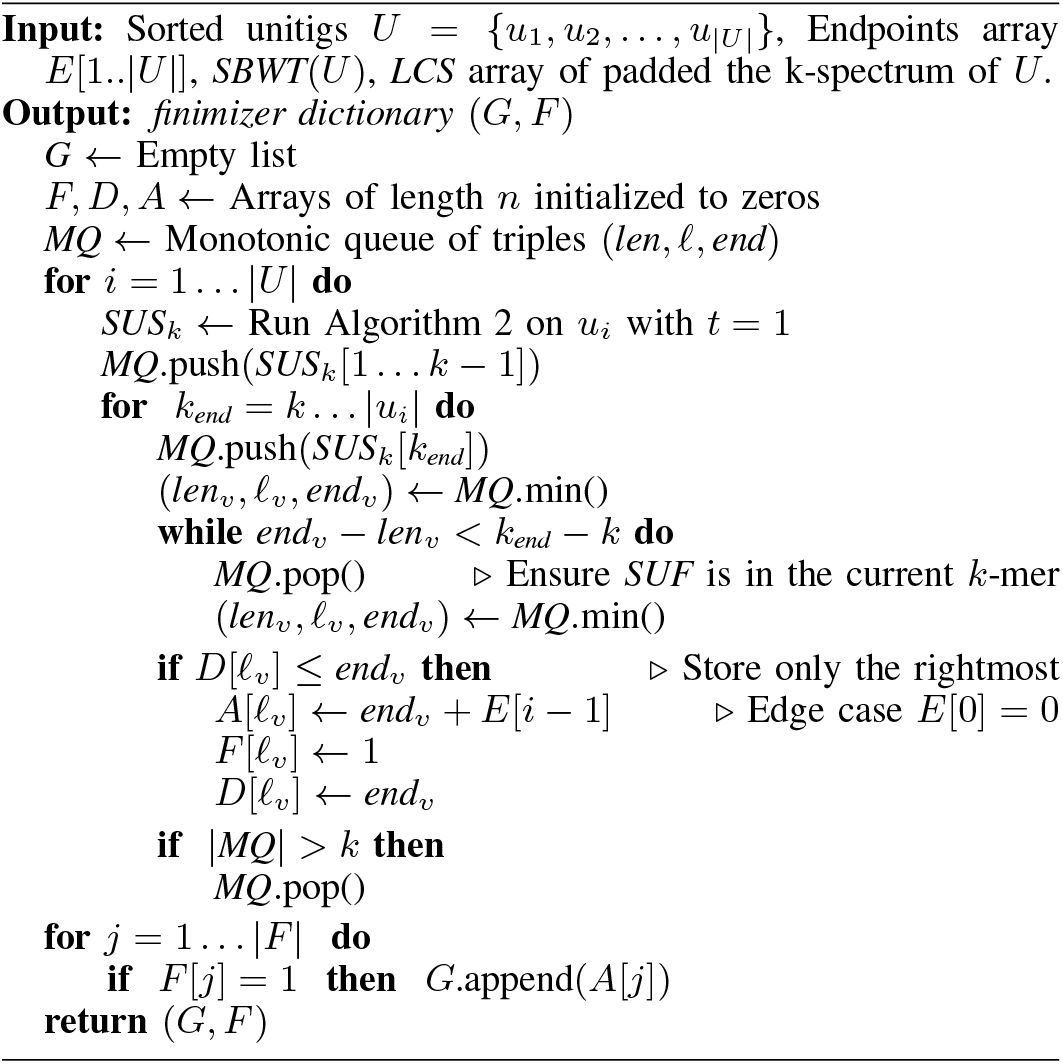

See Figure 1 for an example of Shortest Unique Finimizers and Shortest Unique Suffixes array of a set of unitigs, and Figure 3 for the full index for these finimizers.

### C. Query Algorithm

We are now ready to describe our algorithm for localizing all *k*-mers of a query string *Q* in the set of unitigs, using the shortest unique finimizer index described in the previous section.

The pseudocode is given in Algorithm 4. The first step of the query algorithm involves running Algorithm 1 to obtain *k*-bounded matching statistics in order to identify all *k*-mers of the query that exist in the index. This returns an array, *MS*_*k*_, of size |*Q*|, where each element is a triple (*len, freq, 𝓁*), *len* is the length of the longest match up to that position, *freq* is the frequency of that substring and *𝓁* is the start of its colexicographic interval in the *SBWT*. If *len* = *k*, the whole kmer at position *i* is considered to be found and we will proceed with its localization. As a second step, we run Algorithm 2 and use a sliding window minimum algorithm to pick the finimizer for each *k*-mer that we know exists in the index. This procedure is identical to the one described for identifying *SUF*s in the unitigs to build the finimizer dictionary.

#### Algorithm 4

Streaming Localization Queries

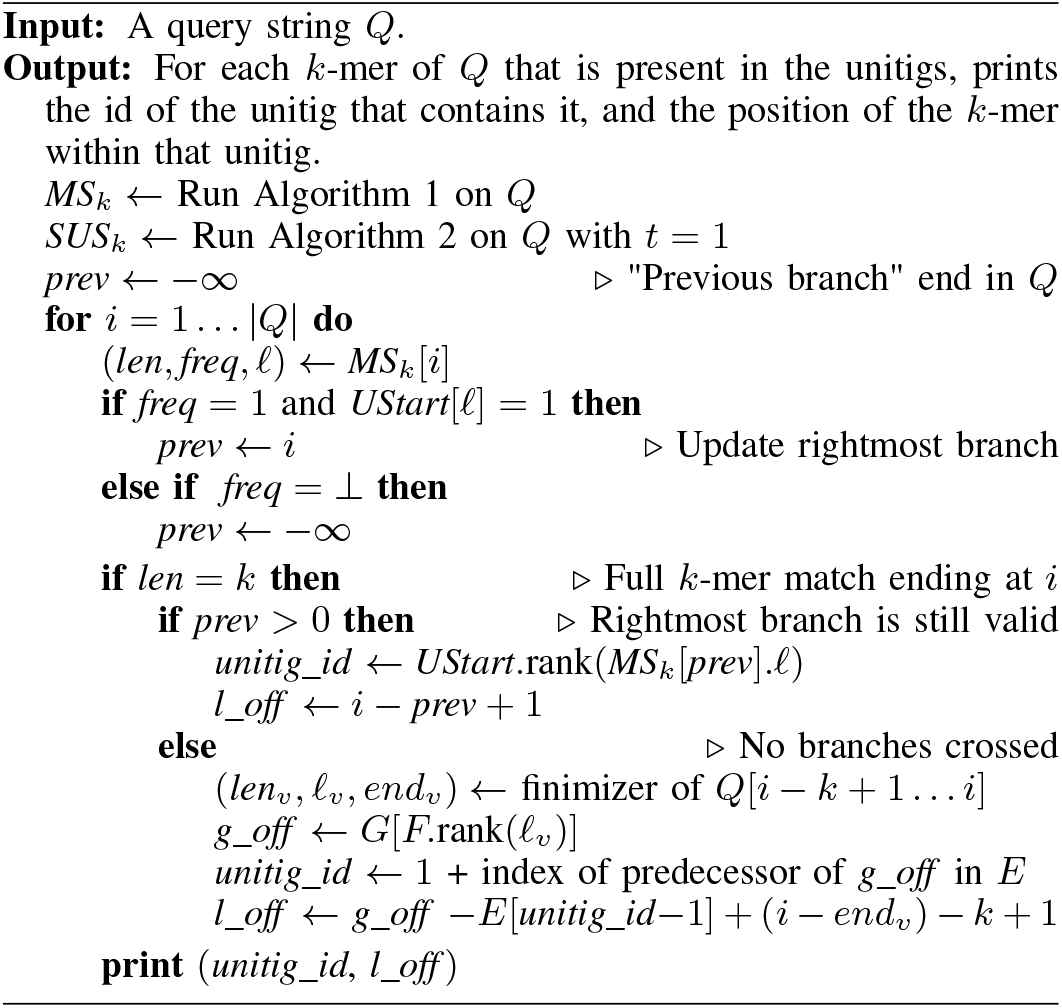

We can locate each found *k*-mer *Y* in the input unitigs in two different ways, using the *branch dictionary* or the *finimizer dictionary*, depending on whether a branch has been observed before the *k*-mer endpoint. For each position *i* of *Q* with *MS*_*k*_[*i*].*freq* = 1 the algorithm checks if the identified unique match is the suffix of the first *k*-mer of a unitig, and if so it assigns the value *i* to *prev*. The starting *k*-mers are marked by the *Ustart* bit vector, thus *prev* is assigned to *i* if we observed a 1 in *UStart* at the colexicographic index stored in *MS*_*k*_[*i*].

As a consequence of this, if a branch has been observed *prev* will have a value bigger than 0, the offset of the closest previous branch. In this case, the unitig identifier of *Y* is obtained, using the *branch dictionary*, with a rank query on *Ustart* since unitigs have been sorted based on the colexicographic rank of their first *k*-mers. The local offset of the *k*-mer *Y* is given by shifting the current position *i* according to the offset of the observed branch, *prev*.

In the case in which no branch has been observed, we use the *finimizer dictionary*. Given the found *SUF υ*, at colexicographic rank *𝓁*_*υ*_, we retrieve, in the global offset array *G*, the global offset (variable *g_off* in the pseudocode) of the unique *k*-mer suffixed by the finimizer corresponding to *𝓁*_*υ*_, which which is at *G*[*F*.rank(*𝓁*_*υ*_)]. The unitig identifier of *Y* can be found with a predecessor query on the unitigs endpoints array *E*.pred(*g_off*). The local offset of the *k*-mer is calculated by subtracting from *g_off* the cumulative length of the previous unitig in the concatenation and shifting the resulting position according to the offset of the finimizer in *Y*.

Examples covering both of the main cases in the localization algorithm can be found in Figure 3. A full proof of the correctness is included in the supplementary material.

### D. Implementation Details

This search algorithm can be improved with what we call a *unitig walk* after each found *k*-mer. This consists of a bitwise comparison of the characters following the found *k*-mer in the query and those in the unitig. For each matching character found, local offset and *k*-mer end, *i*, are augmented by 1. This walk ends when one of the following three conditions is met: 1. the end of the query is reached; 2. the end of the unitig is reached; 3. a mismatch is found. In the latter two cases, *prev* is set back to −∞ and the search continues as described above from the last *k*-mer not yet localized.

It is worth mentioning that in many applications, a *k*-mer is considered equal to its reverse complement. Thus if a *k*-mer *Y* is not found in the *SBWT* of the *DSPSS* it could still be possible to find its reverse complement. We can take care of this by either doubling the query time by querying the *k*-mer in both orientations, or by up to doubling the index size by indexing the reverse complements of the unitigs as well. We can store a permutation mapping the reverse unitigs to their forward counterparts, together with a bit vector marking the starting *k*-mers of the reversed unitigs to easily check the unitig orientation. This allows us to report the locations of the found *k*-mers in the forward unitigs.

## V. Unitig Orientation

Due to the double strandedness of DNA, the existence of a *k*-mer in a genome implies the existence of its reverse complement on the opposite strand. A common space-saving trick is to index only one orientation of each *k*-mer, and consider the other orientation automatically present. For example, unitig extraction tools like BCALM2 [10], CUTTLEFISH2 [23] and GGCAT [11] by default produce only *canonical unitigs*, which are defined to be those unitigs which are lexicographically smaller or equal to their reverse complement. In the best case, this reduces the number of *k*-mers to be indexed by a factor of 2. However, when using the *SBWT* to index the *k*-mers in the unitigs, there is another complication arising from the need to add incoming dummy node paths to all *k*-mers that do not have a predecessor in the de Bruijn graph. If the unitigs are oriented in a canonical way as described above, then due to chance we can expect adjacent unitigs will roughly half the time get oriented in opposite directions, and so the number of unitigs (and therefore *k*-mers) without a predecessor may be unnecessarily large compared to a graph where neighbors are oriented in the same direction. However, since the de Bruijn graph can have cycles, it may be unavoidable that at least some neighbors end up facing in opposite directions, and an interesting combinatorial optimization problem to orient the unitigs emerges.

In this section, we analyze the space overhead caused by the dummy nodes, and describe a heuristic to orient the unitigs to minimize the overhead. A similar heuristic is implemented in Metagraph [22] to reduce the number of dummy nodes in the BOSS data structure. The optimal solution to the problem is left as an open problem.

### A. Dummy node overhead

Let *R* be a set of *k*-mers and *R*^*′*^ ⊆ *R* be the subset of *k*-mers such that *X* ∈ *R*^*′*^ iff ∄*Y* ∈ *R* such that *X*[1..*k* − 1] = *Y* [2..*k*]. Let *D* = _*x*∈*R*_*′* {*x*[1..*i*] | *i* = 1, …, *k* − 1} be the set of distinct proper prefixes of *R*^*′*^. Each element in *D* adds a node to the *SBWT* graph (Figure 1, right). The exact space cost of this depends on the *SBWT* encoding used [3], [4]. For simplicity here we assume each node has a unit cost, and edges are free. Ignoring lower-order terms in the space complexity of rank queries, this model is accurate for the bit matrix encoding of the *SBWT* used for experiments in this paper.

The number of dummy nodes introduced is equal to the number of nodes in the trie of the elements in *D*. Let *x*_1_, … *x*_*n*_ be the list of elements of *R*^*′*^ in lexicographic (note: *not* colexicographic) order. Under the cost model defined above, the total cost is 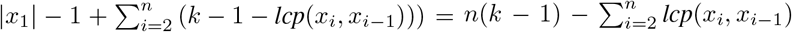, where *lcp*(*x*_*i*_, *x i*−1) is the length of the longest common prefix of *k*-mers *x*_*i*_ and *x*_*i*−1_.

### B. Heuristic

As an exact solution seems difficult, we simplify the problem by just trying to minimize *n* = |*R*^*′*^|. That is, minimizing the number of unitigs without a predecessor. Our heuristic is best described as a series of breadth-first searches on the *bidirected overlap graph* of the unitigs. In a bidirected graph, each endpoint of an edge has an orientation, denoted with ⊕ or ⊖. That is, edges are defined as ordered 4-tuples (*u, v, s*_*u*_, *s*_*υ*_), where *u* and *v* are nodes and *s*_*u*_ and *s*_*v*_ are the orientations of the endpoints. We use these kinds of edges to model all the possible connections between unitigs and their reverse complements. Given a unitig *v*, we denote with *υ*^⊖^ the reverse complement of *v*, and with *v*^⊕^ the unitig *v* itself, and we define that there is an edge (*u, v, s*_*u*_, *s*_*υ*_) iff the last *k* − 1 characters of 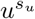 are equal to the first *k* − 1 characters of 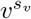. Note that with this definition, the existence of edge (*u, v, s*_*u*_, *s*_*v*_) implies the existence of the symmetric edge (*υ, u*, ¬ *s*_*u*_, ¬ *s*_*υ*_), where ¬ flips the sign of an edge. We call a node *u terminal* iff for every edge (*u, v, s*_*u*_, *s*_*υ*_), the sign *s*_*u*_ is the same.

A breadth-first search maintains a queue of pairs (*u, s*_*u*_), where *u* is a unitig and *s*_*u*_ is the orientation of the unitig. The queue is initialized by pushing an arbitrary terminal node and assigning a forward orientation to it. Then, at each iteration, the queue is popped, and if the node of the obtained pair (*u, s*_*u*_) has not been visited yet with either orientation, we assign orientation *s*_*u*_ to *u*, and push all its neighbors to the queue in orientations in which they overlap with (*u, s*_*u*_). If the queue becomes empty, but not all terminal nodes have been visited so far, we push an arbitary unvisited terminal node with a forward orientation and continue. If all terminal nodes have been visited, we push an arbitrary node. The algorithm concludes when all nodes have been visited. An implementation is available at https://github.com/jnalanko/unitig_flipper.

## VI. Experiments

### Experimental Machine

Our experimental machine had four Intel Xeon E7-4830 v3 CPUs, each running at 2.10 GHz with 12 cores (48 cores total); 30 MiB of L3 cache, and 1.5 TiB of RAM and a 12 TiB serial ATA hard disk. The OS was Linux (Ubuntu 18.04.5 LTS) kernel version 5.4.0-58-generic, with compiler `g++` version 10.3.0 (flags `-O3 -march=native` and `-DNDEBUG`). Runtimes were measured by instrumenting code with calls to the high-resolution clock (`std::chrono`).

### Datasets

We experimented on three different datasets that represent typical targets for *k*-mer indexing (see also Table I).

**TABLE 1.** Statistics on the raw genomic datasets. *k*-mer counts considers canonical *k*-mers. We derived unitigs for each dataset and built a SBWT. $ original is the number of dummy nodes in the original unitig ordering, and $ flipped is the number after applying the unitig orientation heuristic described in Section V.

| | Sequences | Total length | Unique 31-mers | Unitigs | Unitig length | \$ original | \$ flipped |
| --- | --- | --- | --- | --- | --- | --- | --- |
| **E. coli** | 745,409 | 18,957,578,183 | 170,648,610 | 8,939,732 | 438,840,570 | 52,528,157 | 4,838,079 |
| **Metagenome** | 17,336,887 | 8,703,117,274 | 2,761,523,935 | 76,958,596 | 4,182,332,581 | 517,193,809 | 191,843,337 |
| **Nanopore** | 501,216 | 2,888,677,990 | 483,259,051 | 29,581,392 | 1,370,700,811 | 119,248,987 | 10,994,322 |

1) A pangenome of 3682 E. coli genomes^5^.

2) A set of 17,336,887 Illumina HiSeq 2500 reads of length 502 sampled from the human gut (SRA identifier ERR5035349) in a study on bile acid malabsorption [21].

3) A set of 501,216 Nanopore sequencing reads of E. coli genomes (SRA identifier SRR25689478) in a study on assorted bacteria with public health implications^6^.

#### A. Finimizer Length and Density

Figure 4 shows the distribution of lengths of shortest unique finimizers in our three datasets with *k* = 31. The distributions have a peak around length 14 with a long tail to the right. The peak shifts to the right as the number of *k*-mers in the dataset grows. Over 5% of the finimizers in the Nanopore dataset had the longest possible length 31. We think this is due to the high error rate of Nanopore reads: If a true *k*-mer is read by the sequencer at least three times, once without errors, once with an error in the first position and once with an error just before the first position, then neither of its two (*k* − 1)-mers is DBG-unique, and the only possible finimizer is the *k*-mer itself.

**Fig. 4.**
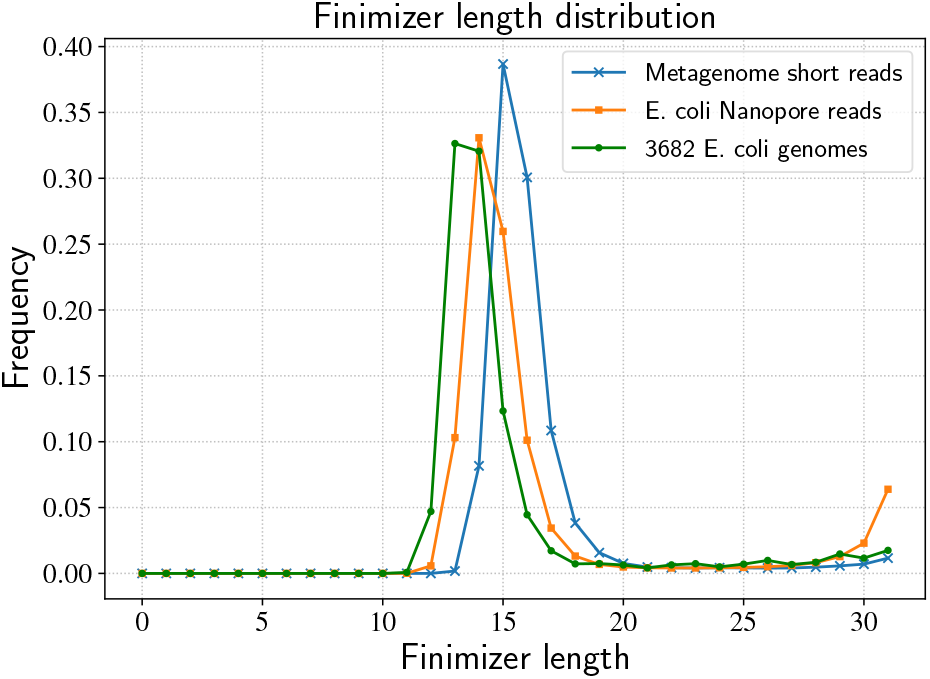
Length distribution of the distinct shortest unique finimizers on three different datasets with *k* = 31.

Figure 5 shows various statistics on shortest *t*-bounded finimizers as a function of *t* in the 3682 E. coli genomes dataset. We find that while the number of distinct finimizers decreases with increasing values of *t*, the total number of DBG occurrences grows. This means that the choice *t* = 1 provides both a smaller index size (since we store a single integer per occurrence) and faster queries as in the worst case the query needs to check all occurrences of a finimizer. However, the average finimizer *length* decreases when *t* grows, which means that indexing strategies whose space depends on the lengths of finimizers (such as those based on the Aho-Corasick automaton) might favor larger choices of *t*.

**Fig. 5.**
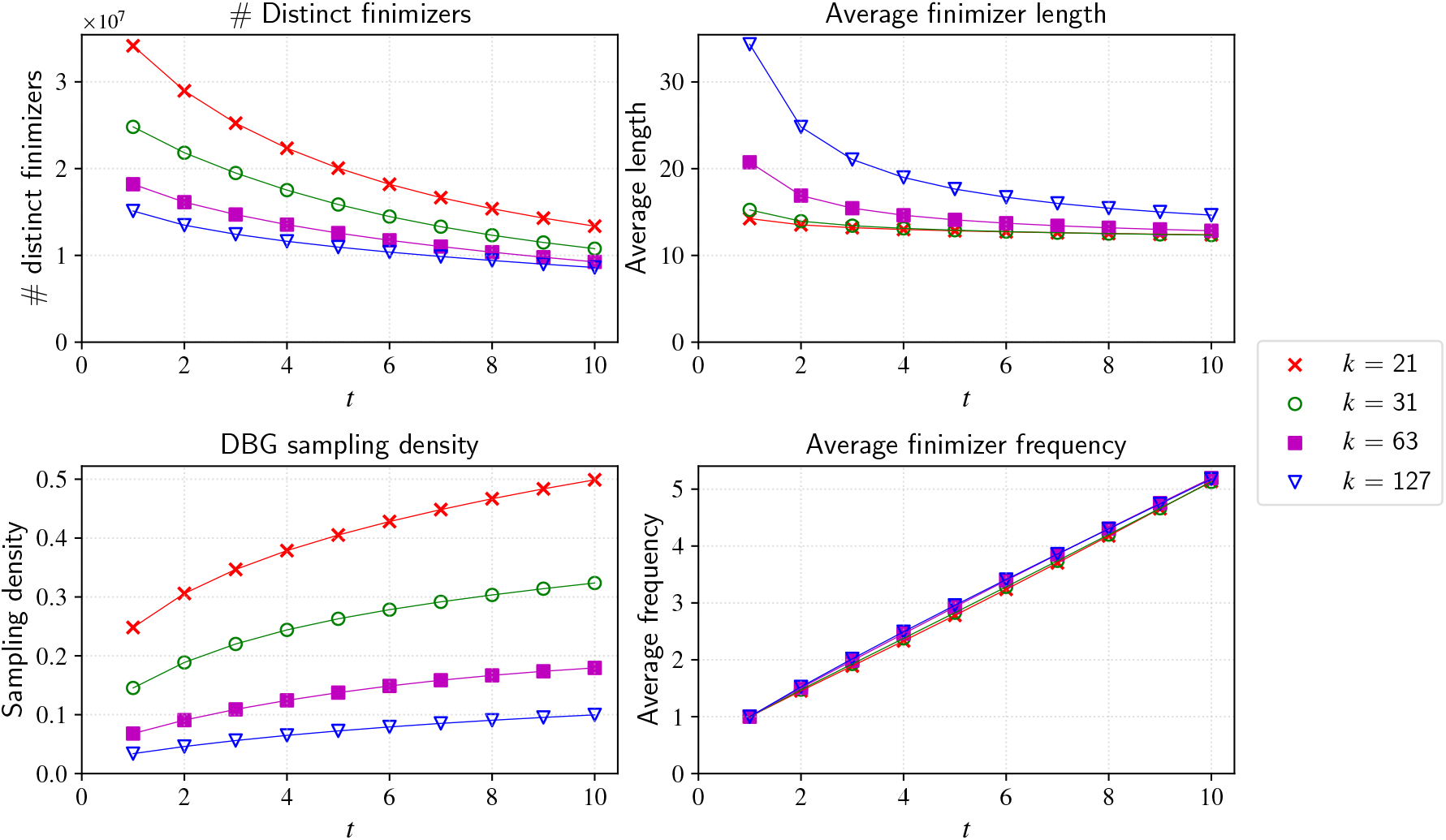
Statistics on shortest *t*-bounded finimizers on the 3682 E. coli genomes dataset. The DBG sampling density at bottom left is defined as the number of DBG occurrences of all distinct finimizers, divided by the number of distinct nodes in the DBG. The frequency plotted at the bottom right is the average number of DBG occurrences of finimizers.

A fixed-length minimizer scheme that orders *m*-mers randomly has a probability of 2*/*(*w* + 1) of selecting a position as the ending position of a minimizer, where *w* = *k* − *m*+1 is the number of *m*-mers inside a *k*-mer [37]. Experimentally, we observed that on random data, this formula also approximately holds for finimizers so that *w* is equal to 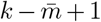, where 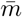 is the average length of chosen finimizers on the dataset: On a random DNA string consisting of 10^6^ characters, with *k* = 63, the average finimizer length was 9.50, and the shortest unique finimizer scheme picked 3.76% positions as ending positions of finimizers, whereas the formula predicts a density of 3.60%. However, on non-random data, the gap becomes wider. On the unitigs of a single E. coli genome (average unitig length 6048) from our 3682 E. coli genomes dataset, 4.3% of positions were selected as ending positions of finimizers, but with average length 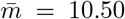, the formula predicts a density of only 3.67%.

#### B. Index Size and Query Performance

We have developed a prototype *k*-mer localization index based on Shortest Unique Finimizers, called FINITO. We now report on performance benchmarks of this prototype on three large datasets. We use SSHASH [33] v3.0.0 (commit 220b2c0) as our performance baseline because it was recently shown to clearly outperform other fast and popular *k*-mer localization tools (e.g., BLIGHT [28])—it is the fastest *k*-mer localization method we know^7^.

### Construction

The input was preprocessed into canonical unitigs before index construction, using GGCAT [11]. The time for this preprocessing in not included in the construction times. We then employed a heuristic method to reduce dummy nodes in the SBWT on all datasets. The method^8^ is described in the supplementary materials, and may be of independent interest.

From here, for *k* = 31 and *m* = 20 SSHASH, with otherwise default parameters, took 11, 3, and 2 minutes to construct, respectively, for the Metagenome, Nanopore, and E.Coli datasets, with peak memory of 11GB, 4GB, and 2GB. For the same *k*, construction of the double-stranded FINITO index was longer, resp.: 299, 58, and 13 minutes (peak memory, resp.: 140GB, 42GB, and 12GB), more than half of which was spent on *SBWT* construction^9^. We remark that although FINITO construction time is higher than SSHASH, it still acceptably fast in real terms. Moreover, the construction of FINITO currently runs on a single thread, whereas SSHASH uses 8. The bottleneck in our construction process—*SBWT* computation—could be significantly sped up with parallelization, a task we leave for future work.

### Queries

We generated two types of query sets: random sequences of length 200 (*negative queries*), the *k*-mers of which are highly unlikely to have any matches in the datasets; and *positive queries* were generated by randomly sampling 200-mers from each dataset. We measured the time FINITO and SSHASH took to look up every *k*-mer in 500,000 positive and negative length-200 sequences, for various values of *k* (see Figure 6). To match Sshash queries, we consider a *k*-mer found if it is found in either forward or reverse complement orientation. As mentioned before, we achieved this either by using an index that contains both the forward and reverse complement strains of all unitigs, and storing the permutation mapping reverse and forward counterparts to report the localizations on the original DNA strand, or by using an index containing only one direction, but running all queries in both directions. The former approach being roughly twice as fast but taking twice the space as the latter. The index size of FINITO was typically larger than Sshash, but the query times of both tools were similar, within a small constant factor from each other, depending on the configuration and the dataset, with negative queries favoring FINITO. Alanko et al. [4] provide several data structures to support *ExtendRight* queries on the SBWT. We use their so-called Plain Matrix representation. Other data structures could reduce the size of our index at the expense of an increased query time, at least with current *SBWT* incarnations. In this respect, it is worth noting that *SBWT* data structures are still very new, and that further engineering may yet lead to significant performance gains.

**Fig. 6.**
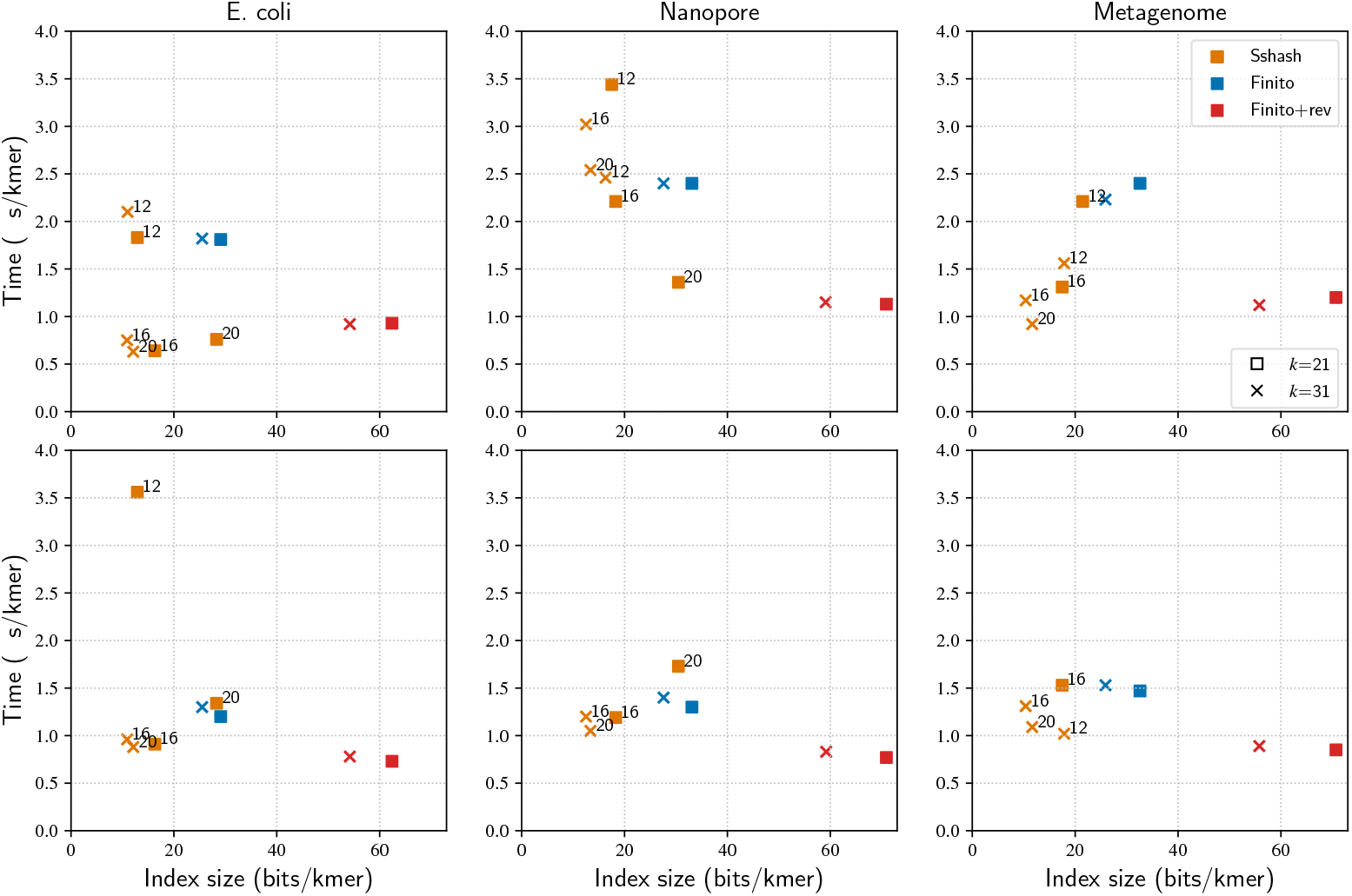
Query performance of SSHASH and FINITO. Top: Time and index size for positive queries for varying *k* and *m* on each dataset. Bottom: Time and index size for negative queries.SShash results for Metagenome with *k* = 21, *m* = 20 are not plotted because construction failed due to collisions of 64-bit hash values.

## VII. Conclusions and Future Work

In the 20 years since minimizers emerged, researchers have invented various heuristics to avoid very frequent minimizers. In this paper we have demonstrated an alternative solution to the problem that works by letting the minimizer length vary. This way, when indexing *DSPSS* representations, we can get a guarantee for the maximum frequency and thus maximum localization query time. A distinct advantage the Shortest Unique Finimizers, we have introduced, have over minimizers is that they are parameter free.

While our current prototype is already competitive with state-of-the-art minimizer-based localization methods, we believe it can be optimized further, e.g., by adding wordlevel parallelism to our *ContractLeft* implementation, and by reducing the time and space for *ExtendRight* by engineering the structures from [4] to use faster recent predecessor [13] and rank [3], [8] structures.

The *SBWT* offers a convenient way to index finimizers for lookup, however more direct methods based on, for example, multiple pattern matching data structures (e.g., [39]), may lead to smaller and faster indexes. The principle challenge (compared to minimizers) is to deal with variable-length keys. In this respect, exploring trie-based solutions may be fruitful.

Finally, interesting applications for finimizers beyond *k*-mer localization include: skipping heuristics in pseudoalignment [6]; approximate *k*-mer matching; and replacing hashing with finimizers in spectrum-preserving tilings [16].

## A Competing interests

No competing interest is declared.

## B Author contributions statement

All authors contributed equally.

## C Acknowledgments

This work was supported in part by the Academy of Finland via grants 339070 and 351150.

## Supplementary material for

## 1 *k*-bounded Matching Statistics

### Algorithm 1

*k*-bounded matching statistics.

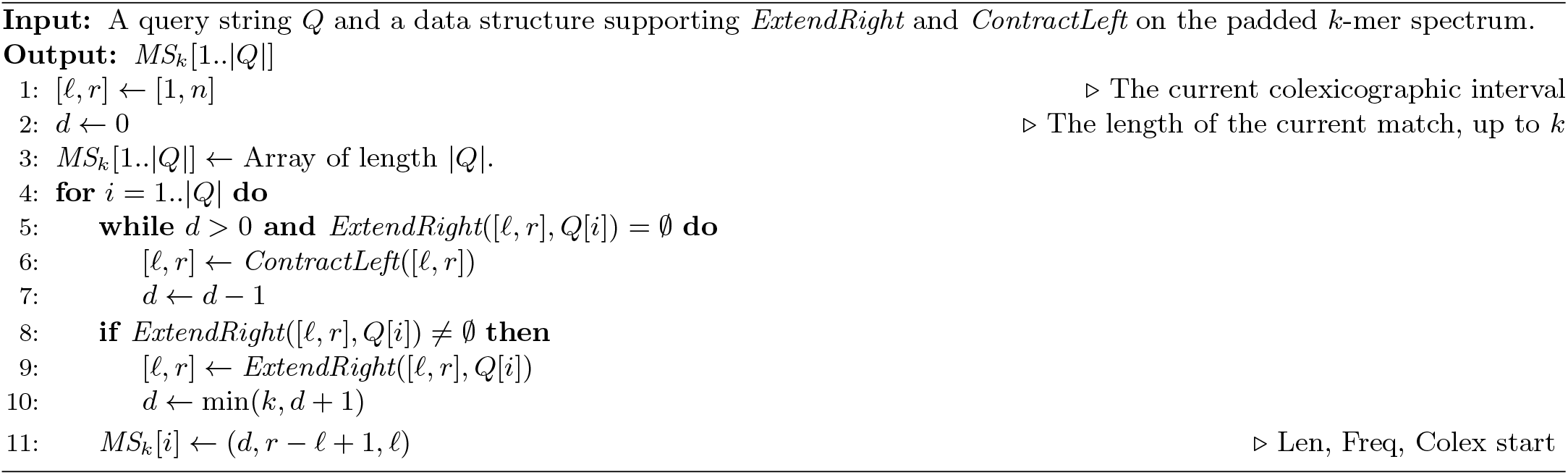

### Theorem 1

*Given constant-time ContractLeft and ExtendRight queries on the padded k-spectrum R*^+^, *Algorithm 1 computes MS*_*k*_ *for query Q in time O*(|*Q*|).

*Proof*. The algorithm maintains the invariant that at the end of iteration *i*, the interval [*𝓁, r*] is the interval of the longest suffix of *Q*[1..*i*] that occurs as suffix of at least one *k*-mer. If [*𝓁, r*] is the longest match at the end of iteration *i* − 1, then at iteration *i* after lines 5 – 7, we have the interval of the longest match ending at *i* − 1 that can be extended with *Q*[*i*], or the interval of the empty string if *Q*[*i*] does not exist in any *k*-mer. If the longest match ending at *i* is nonempty, it is a right extension of such a string, so the invariant is maintained. The algorithm works in *O*(|*Q*|) time because *d* is always non-negative, and the while-loop can run only *d* times because *d* is incremented at most |*Q*| times, and every run of the while-loop decrements *d*.

## 2 Shortest Frequency-bounded Suffixes

### Algorithm 2

Shortest frequency-bounded suffix statistics.

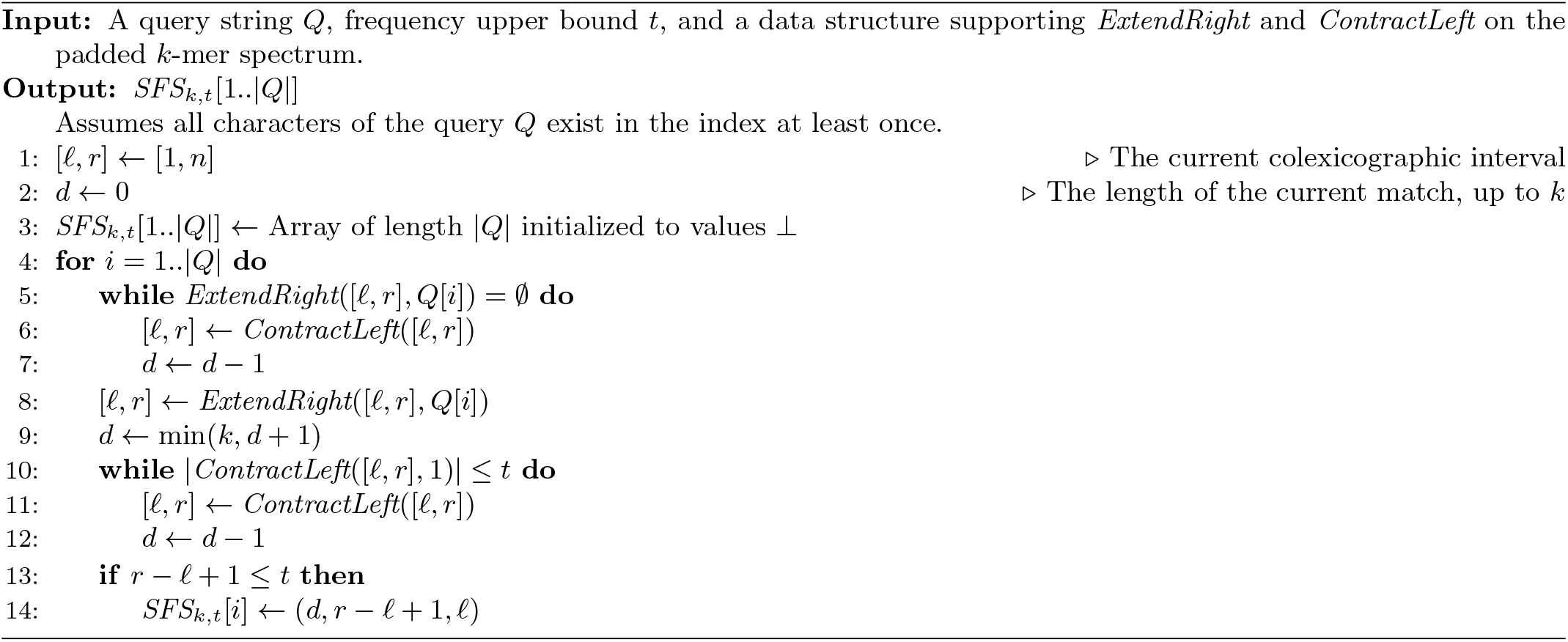

### Definition 2

*(Shortest frequency-bounded suffixes) The shortest suffix statistics of a query sequence Q*[1..*m*], *versus a padded k-spectrum R*^+^, *is the array SFS*_*k,t*_[1..*m*] *such that SFS*_*k,t*_[*i*] = (*d*_*i*_, *p*_*i*_, *𝓁*_*i*_), *where d*_*i*_ *is the smallest positive integer such that Q*(*i* − *d*_*i*_..*i*] *is a suffix of p*_*i*_ *(p*_*i*_ ≤ *t) k-mers in R*^+^, *and 𝓁*_*i*_ *is the start of the colexicographic interval of Q*(*i* − *d*_*i*_..*i*]. *If no such d*_*i*_ *exists, SFS*_*k,t*_[*i*] = ⊥.

### Theorem 2

*Given constant-time ContractLeft and ExtendRight queries on the padded k-spectrum R*^+^, *Algorithm 2 computes SFS*_*k*_ *for query Q in time O*(|*Q*|).

*Proof*. The algorithm maintains the invariant that at the end of iteration *i*, the interval [*𝓁, r*] is the interval of the shortest suffix of *Q*[1..*i*] that occurs as suffix of at most *t k*-mers, if exists, or otherwise [*𝓁, r*] is the longest suffix of *Q*[1..*i*] that occurs in the *k*-mers. To see this, we consider two cases.

- If [*𝓁, r*] is the interval of the longest match at the end of iteration *i* − 1, then in iteration *i*, the lines 5 – 9 update it to the longest match up to position *i*. If the interval becomes of size ≤ *t* after lines 5 – 9, then lines 10 – 12 contract it to the shortest match with frequency at most *t* and the invariant continues to hold. Otherwise if the interval size did not shrink to at most *t* after lines 5 – 9, then lines 10 – 12 are not run and the interval remains the longest match. The invariant continues to hold.
- (ii) If [*𝓁, r*] is the shortest match with frequency at most *t* at the end of iteration *i* − 1, then
  – (iia) if the loop on lines 5 – 7 is run at least once, it becomes the longest match after the right extension at line 8 (a left contraction was needed for a right extension, so the positions that were contracted can not be a part of the longest match). If the loop on lines 10 – 12 is run at least once, it is contracted to the shortest match of frequency at most *t*, otherwise the interval is not changed and it remains the longest match, maintaining the invariant.
  – (iib) if the loop on lines 5 – 7 is not run at least once, then the interval remains unchanged up to line 8, and the right extension on line 8 can only make the frequency smaller, so the frequency remains at most *t*. Lines 10 – 12 contract it to the shortest match with frequency at most *t*, maintaining the invariant.

If the assignment on line 14 happens, it is correct since then [*l, r*] is of size at most *t* and by lines 10 – 12 the length *d* is minimal. If the assignment does not happen, then *SFS*_*k*_[*i*] is correctly left as ⊥ since now the interval [*𝓁, r*] represents the longest available match ending at *i*, and thus a match with frequency *t* or less can not exist, as it would have to be longer than the longest match.

The algorithm works in *O*(|*Q*|) time because the while-loop conditions are false at most |*Q*| times each, and every time they evaluate to true, the left contraction inside can be charged to a distinct right extension on line 8, which is executed |*Q*| times.

## 3 Localization Proof of Correctness

Suppose we have query a *Q* such that *k*-mer *α* = *Q*(*i* − *k*..*i*] **is known to exist** in the DBG according to the matching statistics information. We now prove that our algorithm localizes *α* to the correct unitig.

### Case 1: Localization via the branch dictionary

Suppose there exists a substring *β* = *Q*[*p*..*q*] with *i* − *k* + 1 ≤ *p* ≤ *q* ≤ *i* such that the DBG frequency of *β* is 1, and *β* is the suffix of a *k*-mer that is at the start of a unitig *X*. Further, suppose that *q* is the largest possible (up to *i*) with this property. Then, our algorithm localizes *α* to *X*. Let *δ* = *Q*[*q* + 1..*i*]. First, we claim that *βδ* is a substring of *X*.

− Proof. For a contradiction, suppose this is not the case. Then, since *β* is unique in the DBG and *α* is known to exist in the DBG, it must be the case that *X* ends before the end of *δ*, that is, |*X*| *< k* + |*δ*|, which means that the path labeled with *βδ* contains an edge (*u, v*) such that *u* has outdegree larger than 1. Let *βδ*[1..*r*] be the path label up to and including *v*. This path label is still unique in DBG, and it is a suffix of the first *k*-mer of its unitig. But then *q* was not the largest possible since position *q* + *r* also fulfills the conditions imposed on *q*, contradiction.

Let *α* = *γβδ*. We claim that *γβ* is a substring of *X*.

− Proof. Since *α* exists in the DBG and *β* is unique in the DBG, the occurrence of *β* in the graph has an incoming path with label *γ*. Since *β* = *X*[*k* − |*β*| + 1..*k*], we have that *k* − |*β*| − |*γ*| + 1 ≥ 1 and so *X*[*k* − |*β*| − |*γ*| + 1..*k*] = *γβ*.

Since *β* is DBG-unique, and both *γβ* and *βδ* are in *X*, we conclude that *α* = *γβδ* must also be in *X*.

### Case 2: Localization via the finimizer dictionary

Suppose there does not exist *β* in the case above. Let *ρ* be the shortest unique finimizer of *α*, so *α* = *τρτ*^*′*^. Then our algorithm localizes *α* to the unitig which contains an occurrence of *ρ* the furthest from the start of the unitig. First, we claim that the path of *α* is non-branching DBG.

− Proof. Let *v*_1_, …, *v*_*k*+1_ be the path labeled with *α*. We consider incoming branches and outgoing branches as separate cases:

* Suppose there exists *j >* 1 such that the indegree of *v*_*j*_ is at least 2. Then, *ρ* is the suffix of two distinct *k*-mers, so it is not unique in the DBG, a contradiction.

* Suppose there exists *j < k* + 1 such that the outdegree of *v*_*j*_ is at least 2. Then, *v*_*j*_ must be after the ending point of *ρ* on the path, since otherwise *ρ* is not DBG-unique. But then the label of the subpath from the start of *ρ* to the node *v*_*j*+1_ satisfies the conditions for *β*, which is a contradiction since such *β* was assumed to not exist.

The global offset stored for *ρ* is the one furthest from the unitig start. We claim that the global offset of *ρ* that we have stored is in the unitig that contains *α*:

− Proof. The string *ρτ*^*′*^ is fully contained in the same unitig as *ρ*, because otherwise the path of *ρτ*^*′*^ would be branching, but that cannot be that case since *α* = *τρτ*^*′*^ in non-branching. Since we are at the unitig *X* that contains *ρ* the furthest to the right as possible, the whole string *α* = *τρτ*^*′*^ must be found fully in *X*.

Helsinki Institute for Information Technology (HIIT) Department of Computer Science, University of Helsinki, Finland

## Footnotes

1 Non-branching paths in the de Bruijn graph of the sequences.

2 Our emphasis.

3 SRA identifier SRR25689478, see also Section VI-B.

4 A monotonic queue is a data structure that can report the minimum element in a variable-length sliding window over a stream of data *x*_1_, *x*_2_, … *x*_*n*_. The queue has three operations: Push(*x*): Appends element *x* to the window. Min(): Retrieves the minimum element *x* in the current window along with its index *i* in the window. Pop(): Removes the oldest element in the window. In our use case, the elements are triples of integers (length, frequency, colex rank), ordered lexicographically.

5 The collection is available at zenodo.org/record/6577997.

6 See https://www.ncbi.nlm.nih.gov/bioproject/812595.

7 As described in [33], SSHASH only claims to solve the problem of *k*-mer lookup, but the implementation requires only trival modifications to report *k*-mer localization.

8 See https://github.com/jnalanko/unitig_flipper

9 Construction time for the single-stranded index were faster, resp. 131, 26, 8 minutes (using resp. 4.6, 2, 2.5GB of RAM), with a corresponding slow down in query time.

## Notes

### Competing Interest Statement

The authors have declared no competing interest.

## References

[1] P. Abedin, M. O. Külekci, and S. V. Thankachan. A survey on shortest unique substring queries. Algorithms, 13(9):224, 2020.

[2] J. N. Alanko, E. Biagi, and S. J. Puglisi. Longest common prefix arrays for succinct k-spectra. In Proc. SPIRE, LNCS 14240, pages 1–13. Springer, 2023.

[3] J. N. Alanko, E. Biagi, S. J. Puglisi, and J. Vuohtoniemi. Subset wavelet trees. In Proc. SEA, LIPIcs 265, pages 4:1–4:14. Schloss Dagstuhl, 2023.

[4] J. N. Alanko, S. J. Puglisi, and J. Vuohtoniemi. Small searchable k-spectra via subset rank queries on the spectral Burrows-Wheeler transform. In Proc. ACDA, pages 225–236. SIAM, 2023.

[5] D. Belazzougui and F. Cunial. Indexed matching statistics and shortest unique substrings. In Proc. SPIRE 2014, pages 179–190. Springer, 2014.

[6] N. L. Bray, H. Pimentel, P. Melsted, and L. Pachter. Near-optimal probabilistic RNA-seq quantification. Nature Biotechnology, 34(5):525– 527, 2016.

[7] R. Cánovas and G. Navarro. Practical compressed suffix trees. In P. Festa, editor, Proc. 9th International Symposium Experimental Algorithms (SEA), volume 6049 of Lecture Notes in Computer Science, pages 94–105. Springer, 2010.

[8] M. Ceregini, F. Kurpicz, and R. Venturini. Faster wavelet trees with quad vectors. CoRR, abs/2302.09239, 2023.

[9] R. Chikhi, A. Limasset, S. Jackman, J. T. Simpson, and P. Medvedev. On the representation of de Bruijn graphs. In Proc. RECOMB, LNCS 8394, pages 35–55. Springer, 2014.

[10] R. Chikhi, A. Limasset, and P. Medvedev. Compacting de Bruijn graphs from sequencing data quickly and in low memory. Bioinformatics, 32(12):i201–i208, 06 2016.

[11] A. Cracco and A. Tomescu. Extremely fast construction and querying of compacted and colored de Bruijn graphs with GGCAT. Genome res., 05 2023.

[12] S. Deorowicz, M. Kokot, S. Grabowski, and A. Debudaj-Grabysz. KMC 2: fast and resource-frugal k-mer counting. Bioinformatics, 31(10):1569–1576, 2015.

[13] D. Díaz-Domínguez, S. Dönges, S. J. Puglisi, and L. Salmela. Simple runs-bounded FM-index designs are fast. In Proc. SEA, LIPIcs 265, pages 7:1–7:16. Schloss Dagstuhl, 2023.

[14] B. Ekim, B. Berger, and R. Chikhi. Minimizer-space de bruijn graphs: Whole-genome assembly of long reads in minutes on a personal computer. Cell Systems, 12(10):958–968.e6, 2021.

[15] M. Erbert, S. Rechner, and M. Müller-Hannemann. Gerbil: a fast and memory-efficient k-mer counter with GPU-support. Algorithms for Molecular Biology, 12(9), 2017.

[16] J. Fan, J. Khan, G. E. Pibiri, and R. Patro. Spectrum preserving tilings enable sparse and modular reference indexing. In Proc. RECOMB, LNCS 13976, pages 21–40. Springer, 2023.

[17] U. Ferraro-Petrillo, M. Sorella, G. Cattaneo, R. Giancarlo, and S. E. Rombo. Analyzing big datasets of genomic sequences: fast and scalable collection of k-mer statistics. BMC Bioinformatics, 20(Suppl 4):138, 2019.

[18] J. Fischer. Combined data structure for previous-and next-smaller-values. Theoretical Computer Science, 412(22):2451–2456, 2011.

[19] G. Holley and P. Melsted. Bifrost: highly parallel construction and indexing of colored and compacted de Bruijn graphs. Genome biology, 21(1):1–20, 2020.

[20] C. Jain et al. Weighted minimizer sampling improves long read mapping. Bioinf., 36(Suppl 1):i111–i118, 07 2020.

[21] I. B. Jeffery et al. Differences in fecal microbiomes and metabolomes of people with vs without irritable bowel syndrome and bile acid malabsorption. Gastroenterology, 158(4):1016–1028, 2020.

[22] M. Karasikov, H. Mustafa, D. Danciu, M. Zimmermann, C. Barber, G. Rätsch, and A. Kahles. Metagraph: Indexing and analysing nucleotide archives at petabase-scale. BioRxiv, 2020.

[23] J. Khan, M. Kokot, S. Deorowicz, and R. Patro. Scalable, ultra-fast, and low-memory construction of compacted de Bruijn graphs with Cuttlefish 2. Genome Biology, 23, 09 2022.

[24] H. Li. Aligning sequence reads, clone sequences and assembly contigs with bwa-mem. arXiv preprint 1303.3997, 2013.

[25] H. Li. Minimap2: pairwise alignment for nucleotide sequences. Bioinformatics, 34(18):3094–3100, 05 2018.

[26] C. Marchet, C. Boucher, S. J. Puglisi, P. Medvedev, M. Salson, and R. Chikhi. Data structures based on k-mers for querying large collections of sequencing data sets. Genome Res., 31(1):1–12, 2021.

[27] C. Marchet, Z. Iqbal, D. Gautheret, M. Salson, and R. Chikhi. Reindeer: efficient indexing of k-mer presence and abundance in sequencing datasets. Bioinf., 36:i177–i185, 07 2020.

[28] C. Marchet, M. Kerbiriou, and A. Limasset. BLight: efficient exact associative structure for k-mers. Bioinformatics, 37(18):2858–2865, 2021.

[29] G. Navarro. Compact Data Structures – A practical approach. Cambridge University Press, 2016.

[30] J. Nyström-Persson, G. Keeble-Gagnère, and N. Zawad. Compact and evenly distributed k-mer binning for genomic sequences. Bioinformatics, 37(17):2563–2569, 2021.

[31] E. Ohlebusch, S. Gog, and A. Kügel. Computing matching statistics and maximal exact matches on compressed full-text indexes. In Proc. SPIRE 2010, pages 347–358. Springer, 2010.

[32] Y. Orenstein et al. Designing small universal k-mer hitting sets for improved analysis of high-throughput sequencing. PLoS Computational Biology, 13(10):e1005777, 2017.

[33] G. E. Pibiri. Sparse and skew hashing of K-mers. Bioinformatics, 38(Suppl 1):i185–i194, 06 2022.

[34] A. Rahman and P. Medevedev. Representation of k-mer sets using spectrum-preserving string sets. J. Computational Biology, 28(4):381–394, 2021.

[35] M. Roberts et al. A preprocessor for shotgun assembly of large genomes. J. Computational Biology, 11(4):734–752, 2004.

[36] M. Roberts, W. Hayes, B. R. Hunt, S. M. Mount, and J. A. Yorke. Reducing storage requirements for biological sequence comparison. Bioinformatics, 20(18):3363–3369, 07 2004.

[37] S. Schleimer, D. S. Wilkerson, and A. Aiken. Winnowing: local algorithms for document fingerprinting. In Proceedings of the 2003 ACM SIGMOD international conference on Management of data, pages 76–85, 2003.

[38] S. Schmidt, S. Khan, J. N. Alanko, G. E. Pibiri, and A. I. Tomescu. Matchtigs: minimum plain text representation of k-mer sets. Genome Biology, 24(1):1–32, 2023.

[39] S. Wu and U. Manber. A fast algorithm for multi-pattern searching. Technical Report TR94-17, University of Arizona. Department of Computer Science Tucson, AZ, 1994.

[40] H. Zheng, G. Marçais, and C. Kingsford. Creating and using minimizer sketches in computational genomics. J. Computational Biology, 30(0):1–26, 2023.

